# Prophages of infant-derived *Bifidobacterium longum* subspecies employ antagonistic and synergistic strategies to persist in their host

**DOI:** 10.64898/2026.02.18.706556

**Authors:** James A. D. Docherty, Nataliia Kuzub, Asier Fernández-Pato, Trishla Sinha, Sergio George, Sergio Andreu-Sánchez, Yuvashankar Kavanal, Milla F. Brandao-Gois, Dena Ennis, Lifelines NEXT cohort study, Cyrus A. Mallon, Sanzhima Garmaeva, Moran Yassour, Alexandra Zhernakova

## Abstract

Early colonisation by bifidobacteria is crucial for infant health, with *Bifidobacterium longum* subspecies (*BL.*) dominating the early gut microbiome. However, the interactions between these bacteria and their viruses remain poorly characterised. Here, we applied genomics-based approaches to examine *BL.* prophage composition and dynamics in infants, as well as their antagonistic and mutualistic evolutionary strategies. Across 213 metagenome-assembled genomes recovered from 139 infant faecal samples in the Dutch Lifelines NEXT cohort, 286 previously undescribed prophages were identified and analysed. Comparative genomics revealed extensive viral diversity, evidence of historical recombination, and widespread counter-defence, with ∼80% of prophages encoding anti-CRISPR or CRISPR-evasion proteins. Approximately half of prophages encoded metabolism altering genes. Notably, prophages and host CRISPR spacer arrays were highly stable across longitudinal samples, indicating stable phage‒host associations during early life. Together, these findings show that *BL.* prophages employ antagonistic and synergistic strategies to maintain infectivity and long-term persistence in the infant gut.

## Main

In healthy, full-term infants receiving breastmilk, bifidobacteria dominate the microbiome^1,2^, comprising up to two-thirds of total microbes^3,4^. During breastfeeding, there is substantial mother-to-infant microbial transmission^5^, including of *Bifidobacterium* spp.^6^, with feeding mode acting as one of the strongest influences shaping the early microbiome^7^. Reduced levels of bifidobacteria^8^ and an aberrant microbiota^9,10,11^ during infancy are associated with the future onset of allergic disorders, such as asthma and food allergies, as well as impaired immune development^12^.

In particular, *Bifidobacterium longum* subsp. (*BL.*) *infantis* often establishes itself as the dominant member of the infant gut microbiome due to its conserved ability to consume human milk oligosaccharides (HMOs)^13,14,15,16,17^. *BL. longum* is also a key member of the infant microbiota^18,19^, encoding a more specialised HMO degradation capacity via preferential consumption of fucosylated HMOs^20^. *BL. infantis* and *BL. longum* play key roles in early immunoregulation^21^, maintenance of gut barrier function^22^, and protection from pathogens^23^. Owing to its ability to utilise plant oligosaccharides^14,24^, *BL. longum* can persist through infancy and beyond age six^25^, and is routinely detected in adults^26,27^. While *BL. suis* is predominantly associated with the microbiome of piglets^28,29,30^, recent studies have identified it in human infant faecal samples^31,32^. To date, *Bifidobacterium longum* has six recognised subspecies: *BL. infantis*, *BL. longum*, *BL. suis*, *BL. suillum*, *BL. iuvenis*^29,33,34^, as well as the novel subspecies, *BL. nexti*^35^. *BL. nexti* was recently isolated from stool samples from the Lifelines NEXT (LLNEXT) ^5,36^, Baby Biome Study^37,38^, Dutch Microbiome Project^39^, and MicrobeMom^40^ cohorts, and is extensively described in the accompanying paper (Ennis *et al*., 2026).

A number of studies have now computationally identified diverse prophages in infant- and adult-derived bifidobacterial strains, including from *B. longum*^41,42,43,44^. However, phages of bifidobacteria remain as highly unknown entities, principally because isolation of virulent phages infecting human-associated bifidobacteria has not been successful to date^45,46^. Increasing evidence, however, does implicate *Bifidobacterium* prophages (including *B. longum* subspecies) as key modulators of bifidobacterial populations in the infant gut via induction events^44^. These shifts in bacterial communities can drive dysbiosis because the niche created by a reduction in beneficial bifidobacteria can be reoccupied by potentially harmful microbes via ‘community shufling’^47,48,49^. These antagonistic bifidophage‒host interactions have led to a coevolutionary arms race^41^. Bifidobacteria have accumulated a diverse arsenal of antiviral defence systems, encoding more than double the number of systems than most bacteria on average^41,50,51^, while their infecting prophages have evolved various anti-defences to maintain infectivity^41^.

In this study, we aimed to characterise the composition, dynamics, and genomic diversity of *BL.*-derived prophages. To investigate this, we analysed 213 metagenome-assembled genomes (MAGs) of *BL.* strains obtained from 139 infant samples and assessed their prophage content, longitudinal stability, diversity, and adaptive functions, including anti-CRISPR (Acr) proteins and auxiliary metabolic genes (AMGs). This allowed us to identify specific antagonistic and mutualistic strategies employed by *BL.* prophages to maintain their infectivity and persistence.

## Results

### *Bifidobacterium longum* subspecies host genomically diverse prophages

To characterise the *BL.* prophage landscape, we analysed 322 available MAGs from LLNEXT infant samples (see Methods). After strain-level dereplication, the final database of *BL.* strains included 213 MAGs from 139 infants from 137 families, comprising *BL. infantis* (n = 123), *BL. longum* (n = 53), *BL. nexti* (n = 31), and *BL. suis* (n = 6). These 213 *BL.* MAGs had a median GC content of ∼60% and a median genome size of ∼2.5 Mbp. GC content showed very little variation across strains and subspecies (SD = 0.24), although there were significant differences in genome size (p ≤ 0.05, Kruskal-Wallis test) (Figure S1). When comparing individual subspecies, *BL. infantis* (2.61 Mbp) had a significantly larger genome size than *BL. longum* (2.35 Mbp), *BL. nexti* (2.37 Mbp) and *BL. suis* (2.32 Mbp) (p ≤ 0.05, Dunn’s test), consistent with previous work based on isolate analysis^35^. All metadata associated with the *BL.* MAGs is provided in Supplementary Table 1.

To understand the prophage content of *B. longum* subspecies, they were mined for putative proviral sequences. After strict quality control (see Methods), this left us with a complete database of 286 ‘high-quality’ prophages from 150 of the 213 (∼70%) MAGs. Analysis of prophage content between individual subspecies revealed a significant association between host subspecies and prophage carriage (p-value = 0.009, χ² = 11.65, df = 3, Pearson’s chi-square test of independence). At least one prophage was detected in ∼73% of *BL. infantis* (90/123) and ∼77% of *BL. longum* (41/53) strains and in all but one *BL. suis* strain (5/6; ∼83), but in just ∼45% of *BL. nexti* strains (14/31) (Figure S2). While the proportion of prophage-containing strains in *BL. infantis*, *BL. longum*, and *BL. suis* (∼75%) was consistent with previous reports^41^, the abundance of prophages in the newly identified subspecies *BL. nexti* was markedly lower. In all, we identified 190 prophage from *BL. infantis*, 72 from *BL. longum*, 17 from *BL. nexus,* and 7 from *BL. suis*. Together, these data demonstrate that infant-derived *BL.* strains have been successfully infected by temperate phages, indicating the presence of proviral populations in the infant gut that have evolved mechanisms to circumvent host defences. All prophage-associated metadata is included in Supplementary Table 2.

We investigated the taxonomy of the recovered prophages, and all were annotated as previously undescribed *Caudoviricetes* phages. We assigned these prophages to *de novo* taxa, comprising 12 families, 69 genera and 80 species, providing early indications of high genomic diversity within this dataset of prophages. Within each species-level cluster, we selected one representative prophage sequence based on it having the largest genome size within the group. The genome length of prophages ranged from 9,174 bp to 79,935 bp, with a median length of 32,781 bp. The GC content of prophages (∼62%) was significantly higher than that of their bacterial hosts across 150 host genomes (∼60%) (p = 2.2e^-16^, V = 11,013, Wilcoxon signed-rank test on host-level means). We compared intergenomic similarity (%) across all prophage sequences to explore genomic diversity, evidence of historical recombination, mobility, and pervasiveness (Figure 1). While there were several distinct regions of clustering, prophages demonstrated extensive genetic diversity, with most sharing <70% pairwise nucleotide identity. Prophages also appeared to cluster predominantly by host subspecies rather than phenotypic factors, such as birth- and feeding mode, with groups of similar prophages predominantly infecting specific host subspecies. In some distantly related prophages, we did observe areas of intermediate intergenomic similarity (∼30‒50%) (see the peripheral regions of the heatmap in Figure 1), suggesting possible historical recombination of viral genomes.

**Figure 1.**
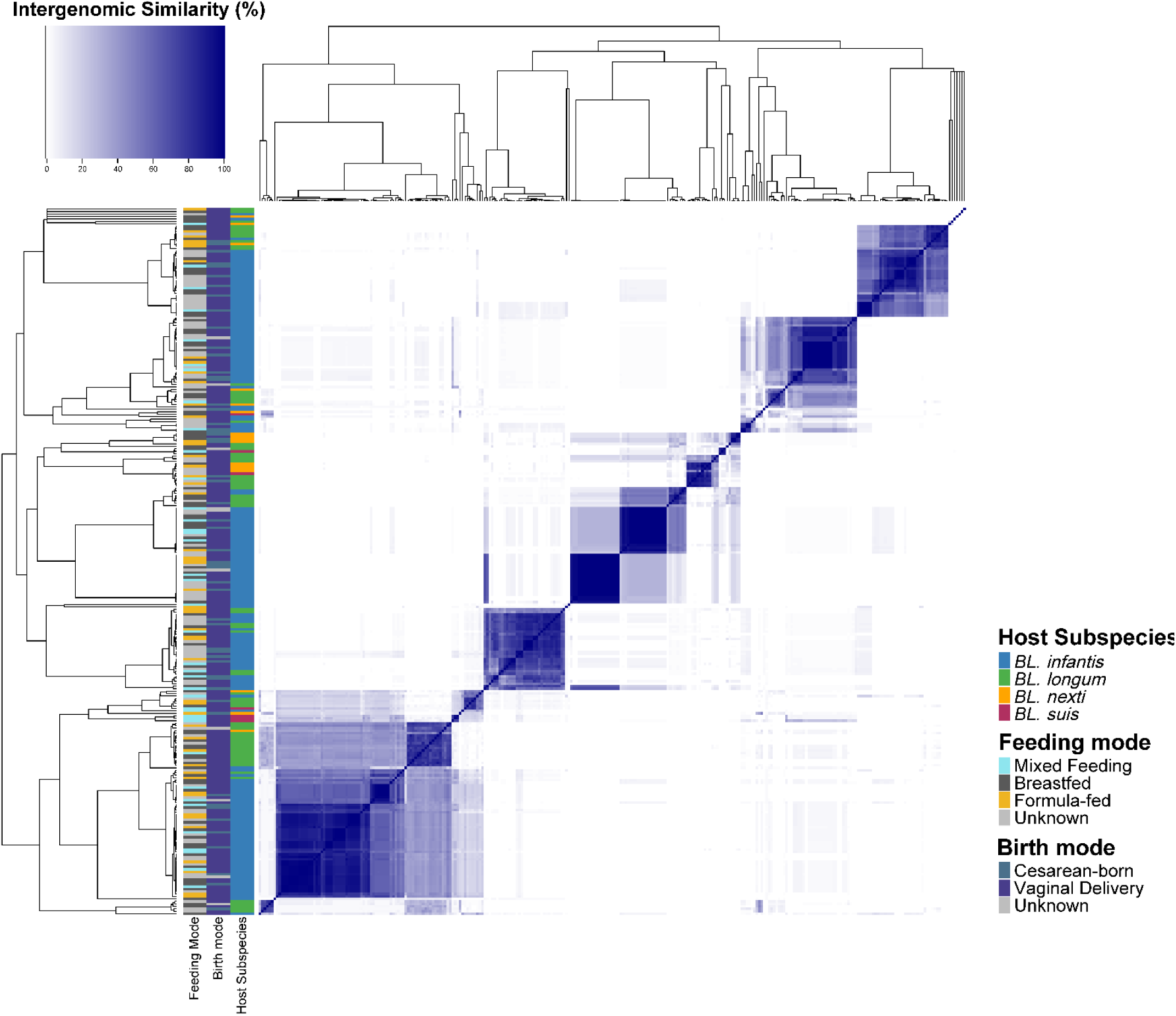
Pairwise comparison of percentage (%) intergenomic similarity (heatmap) and hierarchical clustering (dendrogram) of 286 *BL.*-derived prophage genomes. Proviral genomes cluster in numerous distinct groups based on intergenomic similarity. Colour strips at left of the heatmap correspond to phenotypic metadata: host subspecies, infant birth mode and infant feeding mode.

After intergenomic comparisons of prophages, their evolutionary relationships were explored by generating a proteomic tree derived from genome-wide amino acid comparisons (Figure 2a). As the phages initially separated into three major clades, this likely represents a minimum of three viral orders. Within these groups, prophages clustered into numerous subclades of varying branch lengths, with shorter branch lengths signifying recent divergence and deeper branching clades indicating more unique evolutionary histories. We again saw that most closely related phages appeared to be confined to a single host (Figure 2a). Interestingly, three distinct clades of near identical prophages infect at least 10 *BL. infantis* strains, highlighting the presence of prophages with widespread distributions across this particular subspecies. Two of these prophages had a relatively small genome size of ∼19kbp (Figures S3a,b, Figure 2a, yellow and green circles), while the third ‘widespread’ *BL. infantis* prophage had a larger genome length of ∼40 kbp (Figure S3c, Figure 2a, dark blue circle). Each of these prophages encode a gene assigned to the ‘moron, auxiliary metabolic gene and host takeover’ (Mo-AMG-HT) PHROG category (Figures S3a–c). Genes within this category are typically horizontally acquired and non-essential, rather they are thought to modulate host metabolism or physiology in ways that enhance viral replication^52,53^. The presence of Mo-AMG-HT genes among widespread *BL. infantis* prophages suggests they may contribute to host adaptation and long-term prophage maintenance and tolerance. Overall, the presence of genomically diverse prophages with distinct evolutionary trajectories indicates several different evolution patterns for maintaining infectivity and persistence within *BL.* populations.

**Figure 2.**
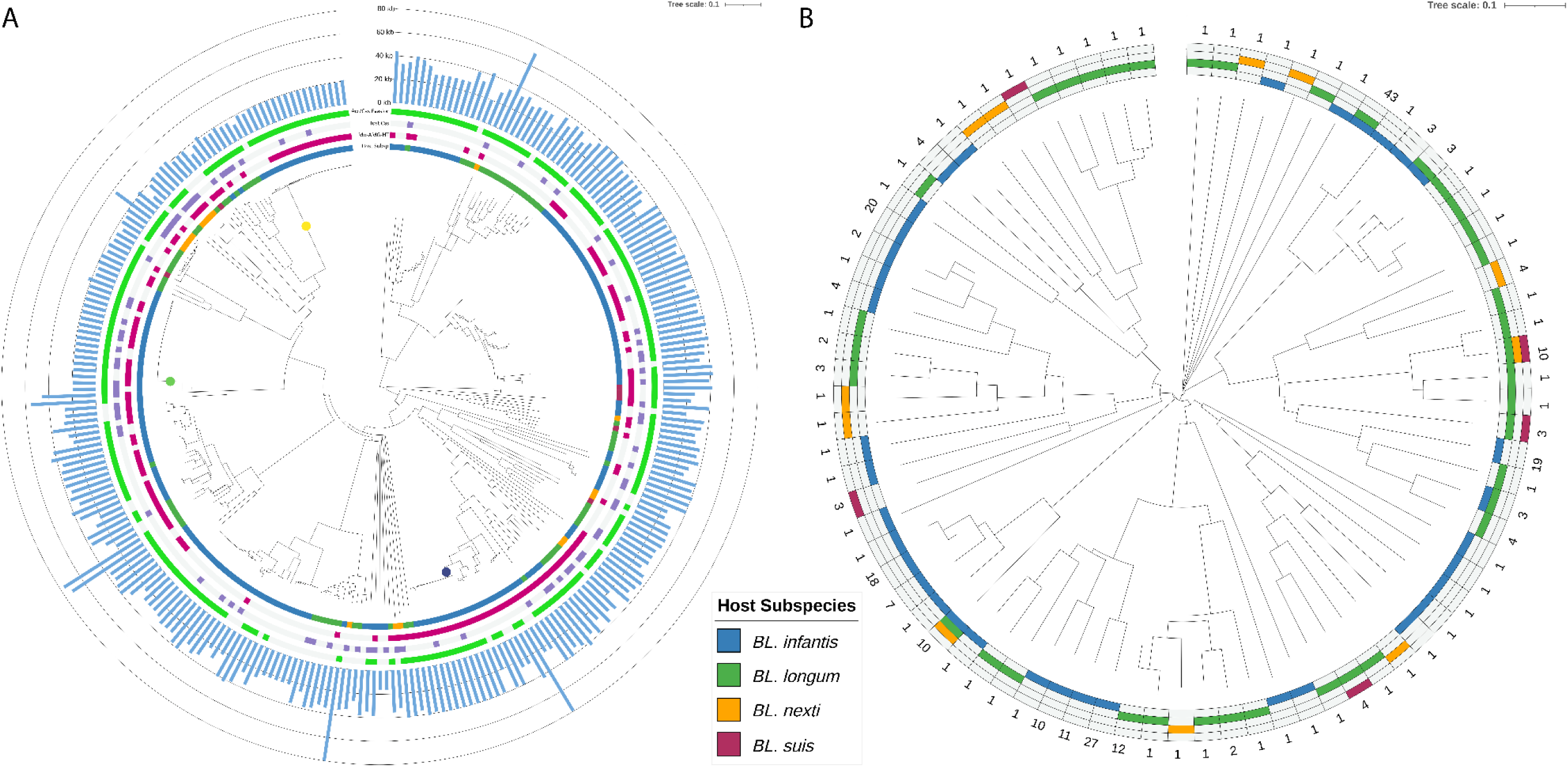
Whole-proteome phylogenetic distance trees of *BL.*-derived prophages. **a**, Whole-proteome phylogenetic tree of all 286 *BL.*-derived prophages. From the inner to outer ring: host subspecies in which each prophage was identified (Host Subsp), presence of genes from the ‘moron, auxiliary metabolic gene and host takeover’ PHROG category (Mo-AMG-HT), presence of a host CRISPR-Cas system (Host Cas), presence of an anti-CRISPR (Acr) or a protein associated with CRISPR-Cas evasion via DNA repair (Acr/Cas Evasion), and prophage genome length. **b**, Whole-proteome phylogenetic tree of 80 viral species representatives. The surrounding heatmap indicates the host subspecies in which each species was found. The numbers denote the number of prophage genomes belonging to each species.

### *BL.*-derived prophages exhibit mobilisation and mechanisms for host-range expansion

To explore the prophage mobilisation patterns and further examine phylogenetic relationships, we constructed a proteomic tree with the 80 viral species representatives, including the number of host subspecies present within each proviral species (Figure 2b). Just 25/80 (∼31%) viral species identified had more than one prophage within the cluster, further highlighting their genomic diversity. Additionally, only seven species had members infecting two or more host subspecies. While overall mobility was limited, two viral species had members detected in three host subspecies, indicating the presence of either broad host range phages or the occurrence of host-switching events. To identify the genetic features that may have enabled host-range expansion, we performed gene cluster comparisons using pairwise alignments of translated proteins across prophage genomes (Figure 3a). This revealed several completely unique and non-homologous genes in prophages of the same viral species, revealing clear evidence of previous horizontal gene acquisition. These distinct genes encoded functions for integration and excision, transcriptional regulation, and nucleotide metabolism, as well as multiple proteins of unknown function. As integrases are dependent on highly specific target sequences^54^, and thus often limited to a narrow host range^55^, the acquisition of a novel integrase could enable the prophage to integrate into a new host with a reciprocal attachment site. Additionally, incompatibilities at transcription and replication stages cause phage infections to be aborted^56^, suggesting that the acquisition of novel regulatory or metabolic genes may facilitate adaptation to new host environments.

**Figure 3.**
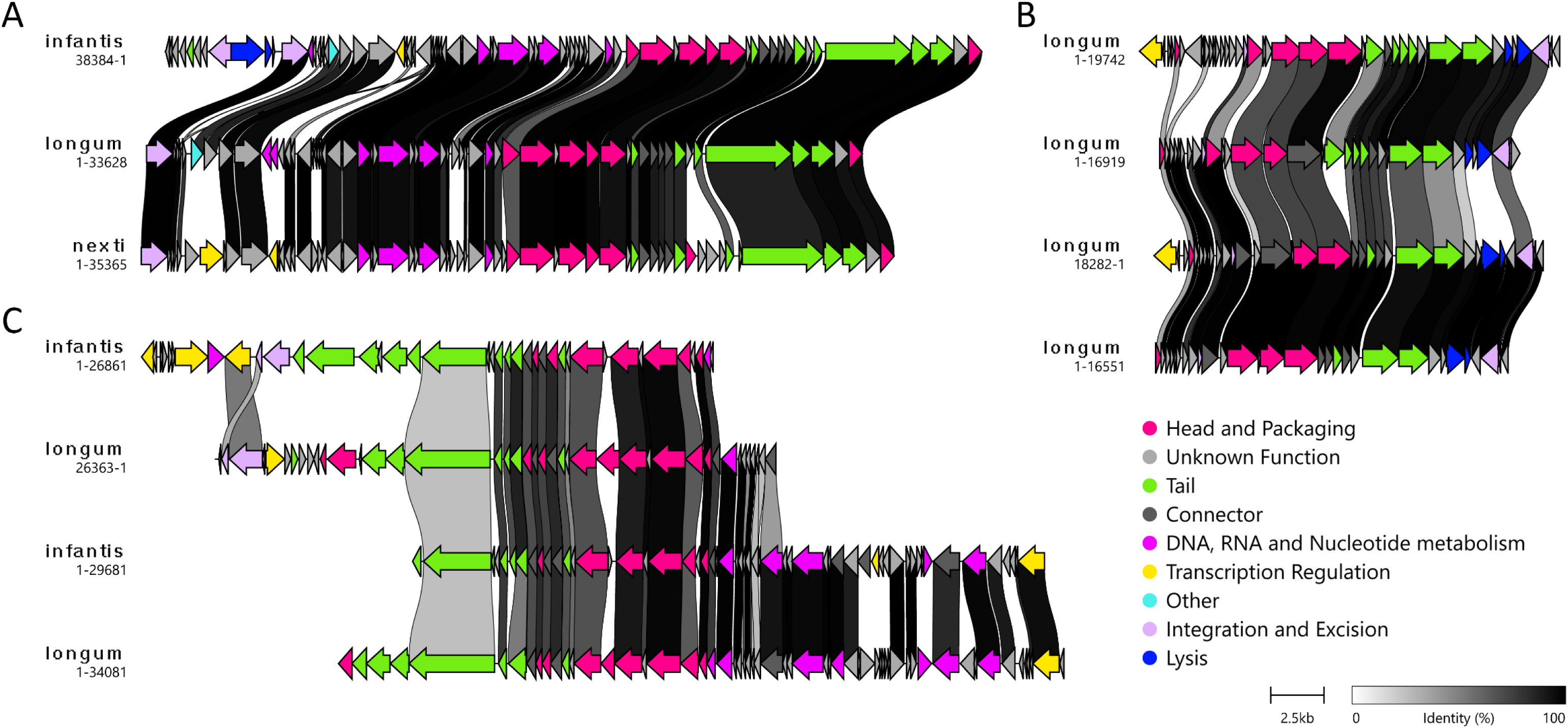
Gene cluster comparisons of *BL.*-derived prophages. **a**, Gene cluster comparison showing evidence of horizontal gene acquisition within a prophage species whose members infect three different *B. longum* subspecies. **b**, Prophages with highly similar overall genome organisation but clear accumulation of mutations across shared modules, all infecting *BL. longum*. **c**, Certain prophage genomes displayed strong conservation in capsid genes but little to no similarity across other regions, including tail cassettes, which is consistent with historical in-host recombination and modular genome replacement.

We also explored alternative mechanisms for host-switching or host-range expansion. First, we performed gene cluster comparisons between prophages with similar genetic architectures that infect multiple members of a single host subspecies (Figure 3b). Several prophages infecting *BL. longum* appeared to have accumulated genomic mutations in multiple structural genes, including genes coding for assembly of the capsid and tail, as well as DNA packaging genes. Consequently, the accumulation of mutations in essential genes, such as receptor-binding proteins of the tail or tail-fibres, could have altered the specificity of these prophages^57,58^. Further probing of mutation-driving mechanisms revealed that 26/286 (∼9%) prophages had tail-targeting diversity-generating retroelements (DGRs), a rapid mechanism for introducing mutations to alter phage host range^59,60,61^. Given that recombination is also known to drive phage host-range expansion^62^, we compared gene clusters from distantly related prophages infecting different hosts, but sharing intermediate intergenomic similarity, in order to identify evidence of historical genetic recombination (Figure 3c). Here we observed capsid proteins with complete homology in each prophage but extensive divergence across the cassette of tail genes. These patterns are highly indicative of a historical modular recombination event in which these capsid gene modules were exchanged between prophages. While we observed only moderate similarity between each tail length tape measure protein, its consistent presence indicates that it was simultaneously recombined with the capsid genes but has undergone progressive accumulation of mutations since the mobilisation event.

Collectively, these results suggest that host-switching and expansion, enabled by multiple distinct strategies, contribute to the long-term persistence of *BL.*-derived prophages, even as host resistance evolves.

### *BL.* strains encode phage-targeting CRISPR-Cas systems

Based on the known antagonistic co-evolution and accumulation of defence and anti-defence systems in human-associated bifidobacteria and prophages^41^, we first looked for phage-targeting CRISPR-Cas systems in host bacterial genomes (see Methods). Collectively, CRISPR-Cas systems were identified in ∼46% (98/213) of all *BL.* strains. Though, large disparities were observed in CRISPR-Cas prevalence across subspecies, with Cas systems detected in ∼94% of *BL. nexti* strains (29/31), compared with ∼38% of *BL. infantis*, *BL. longum*, and *BL. suis* strains. While the presence of CRISPR-Cas systems in these three subspecies was consistent with previous reports (∼38%)^63^, the near-universal presence of Cas systems in *BL. nexti* highlights a steep contrast in their metabolic dedication to adaptive defence. Again, a clear disparity was observed in trends between *BL. nexti* and the other *BL.* strains, with *BL. nexti* harbouring much higher numbers of CRISPR-Cas systems and significantly lower numbers of prophages. Also similar to previous findings^63,64^, Cas subtypes I-C (n = 42), I-E (n = 48), I-G (n = 1), and II-C (n = 21) were detected in this dataset of *BL.* strains (Figure S4). Notably, multiple strains encoded two distinct Cas subtypes simultaneously, further underscoring the broad antiviral defence capabilities of several *BL.* strains (Figure S5).

To confirm that the targets of Cas systems were viral, we compared CRISPR spacer sequences from host strains against *BL.*-derived prophages to identify homologous matches. This identified 6,304 spacer sequences in ∼61% (130/213) of bacterial strains (median = 39 spacers per strain). As complete CRISPR-Cas systems were found in 98 strains, 32 strains likely possess orphan arrays or degraded CRISPR-Cas systems. In the 6,304 non-dereplicated spacer sequences, there were 2,800 spacer hits targeting 135/286 (∼47%) *BL.*-derived prophages. The largest portion of phage-targeting sequences were for genes encoding unknown functions (952/2,800, 34%), highlighting the extensive proviral dark matter present within bifidobacteria. However, when spacer hits targeting structural proteins were combined (n = 1,039), including head and packaging (n = 486), tail (n = 444), and connector (n = 109) genes, they collectively represented the most frequently targeted category of phage genes by CRISPR spacers. There were also spacers that shared homology with viral genes belonging to each of the remaining six PHROG categories (n = 822) (Figure 4a). Another 96 spacers matched non-coding sequences of proviral genomes. Dereplication of spacers at 100% sequence homology revealed 1,943 non-redundant spacers, indicating a large number shared spacers between strains. This could reflect horizontal transfer of CRISPR cassettes between strains, a potential mechanism for inherited prophage resistance^65^. The diversity of CRISPR-Cas types observed, coupled with the high abundance of confirmed prophage-targeting CRISPR spacers, suggests that *BL.* strains acquire adaptive resistance to invasive proviral elements. Furthermore, the targeting of both essential and accessory phage genes indicates that these systems have diverse targets for conferring resistance.

**Figure 4.**
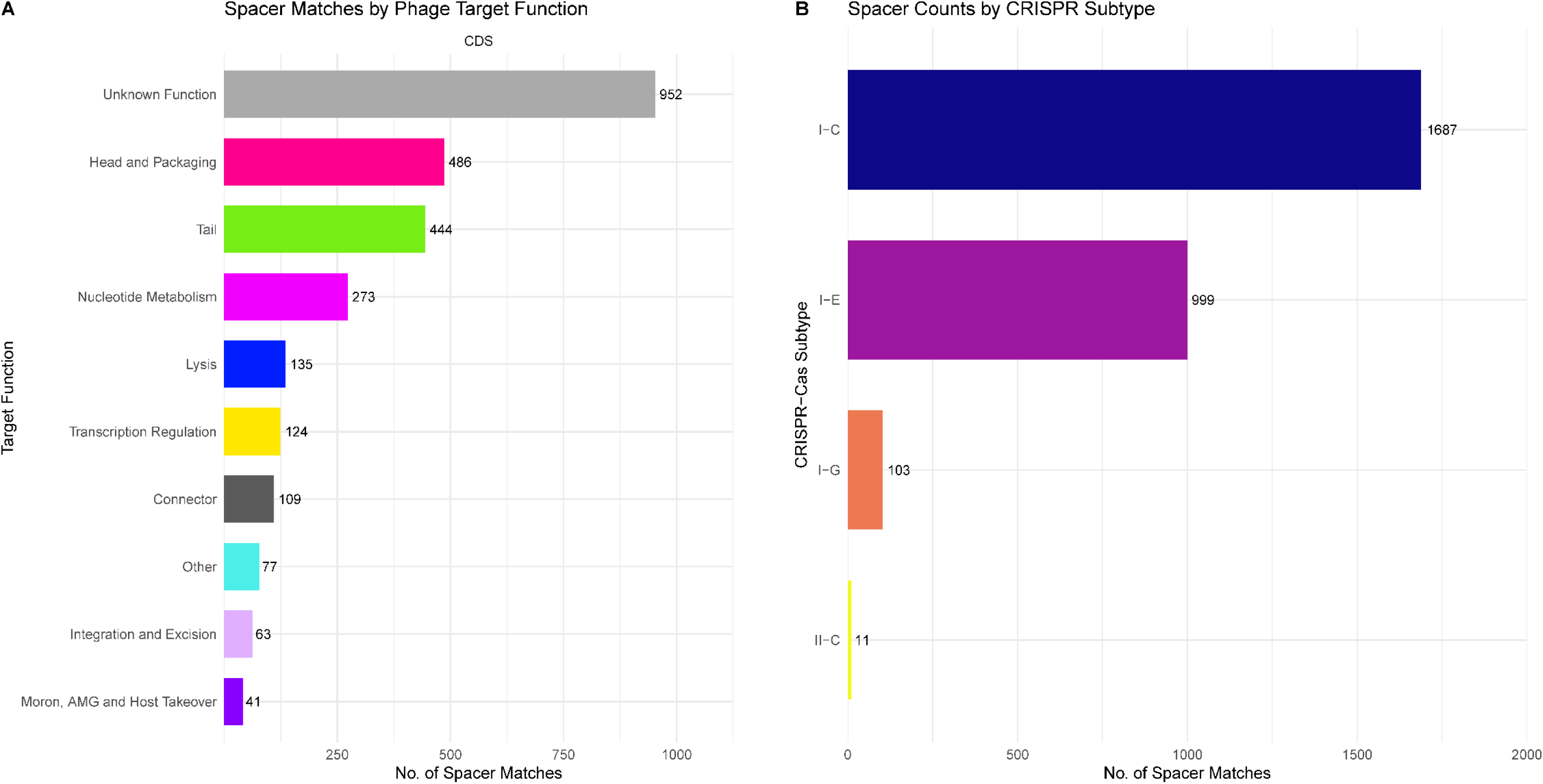
CRISPR spacer targeting of prophages detected in *BL.* strains. **a,** Functional categories of viral genes targeted by spacers based on PHROG annotations. Bars show the total number of spacer hits per functional category. **b**, Distribution of spacer matches by CRISPR-Cas subtype.

Examination of the origin of phage-targeting spacers revealed that 2,789/2,800 spacer‒ phage matches (99.6%) originated from type I systems (Figure 4b), in agreement with previous research on human-associated bifidobacteria^41^. Spacers belonging to CRISPR-Cas type I-C accounted for 1,687 spacer matches, and types I-E and I-G accounted for 999 and 103 hits, respectively. Extending upon previous findings, these results suggest that it is type I CRISPR-Cas systems that primarily confer resistance to phages in bifidobacteria, rather than other invasive mobile genetic elements (MGEs). Of the 21 Type II-C systems detected in *BL.* strains, only 11 spacers from four genomes exhibited homology to prophage sequences, suggesting that these systems primarily target other MGEs, such as plasmids. Additionally, only six strains carried spacers targeting prophages within their own genome, consistent with negative selection driving the removal of spacers that could trigger autoimmune fitness costs^66^. Among the three widespread prophages that infect 10 or more *BL. infantis* strains, only one was targeted by three CRISPR spacers, suggesting extensive domestication and tolerance by their host strains. Further examination of spacer targeting patterns revealed that *BL. nexti* strains accounted for ∼32% (906/2,800) of all spacer‒phage matches, even though they comprised only ∼15% of all MAGs. This disproportionately high contribution of phage-targeting spacers, together with the prevalence of CRISPR-Cas systems in *BL. nexti*, may help explain the significantly lower abundance of proviral sequences observed in this subspecies.

### *BL.* prophages encode various anti-defence systems

Given our identification of CRISPR-Cas systems with spacers mapping to the prophages we detected, we next examined counter-adaptive immunity mechanisms to determine how these proviral elements circumvent host defences. First, we explored the presence of anti-CRISPR (Acr) proteins, identifying 223 Acrs in 147/286 (∼51%) prophages. Of the 24 distinct Acr types detected, five were found in 20 or more prophages. We then investigated these five more closely. In line with the identified CRISPR-Cas systems, the Acrs found here were collectively predicted to inhibit CRISPR types I-C, I-E, and II-C^67^, despite not having yet been experimentally validated. As four of these five Acr proteins had at least 10 non-identical amino acid sequences, we selected them for tests of selection pressure. Based on the Nei-Gojobori method^68^, three of these Acrs showed significant evidence of codon-based purifying selection (p < 0.05, Fisher’s exact test), implying strong functional conservation and limited genetic diversification^69^. This is consistent with an essential role for these proteins in prophage maintenance. As it is also thought that purifying selection predominantly acts on highly expressed genes^70^, these results are also consistent with ongoing utilisation of these Acrs within host cells.

We next investigated alternative non-Acr anti-defence systems, which revealed 365 individual systems present within 188/286 (∼66%) prophages, spanning 10 different classes/types. The most common non-Acr anti-defence system was ‘CRISPR-Cas evasion by DNA repair’, with 156 individual occurrences in 146 prophages. When combined with Acr systems, 233/286 (∼81%) of *BL.*-derived prophages possess either a CRISPR-evasion protein or an Acr, with 147 phages encoding both. There were also anti-defence mechanisms to circumvent the host SOS response (n = 110), restriction modification (RM) (n = 47), RecBCD enzymes (n = 16), and Gabija (n = 4). A single counter-defence mechanism was identified across several prophages, including Antiviral STAND (n = 1), bacteriophage exclusion (n = 1), Dnd (n = 1), and toxin-antitoxin (n = 1). Finally, we detected 28 individual proteins with broad-spectrum counter-defence functions.

In summary, the presence of conserved Acr proteins and CRISPR-Cas alleviation mechanisms in >80% of prophages demonstrates that they have evolved specialised and specific mechanisms to circumvent the adaptive defences of their host. Additionally, the presence of anti-defences targeting innate immune mechanisms, such as anti-RM, show their anti-defence repertoire is not confined to inhibiting CRISPR-Cas systems. However, our identification of larger numbers of Acrs and CRISPR-Cas evasion proteins relative to other anti-defence mechanisms likely reflects a discovery bias, as Acrs are currently far better characterised and represented in databases^71^. In contrast, many non-Acr defence-evasion strategies remain poorly described^72^, suggesting that additional systems exist but are not yet detectable using current computational tools.

### Prophages from *BL.* strains possess multiple genes for altering host metabolism

We investigated additional potential antagonistic mechanisms underlying phage persistence by looking for viral genes capable of modulating or ‘hijacking’ host metabolism. Putative mutualistic strategies were assessed in parallel by identifying prophage-encoded antibiotic resistance genes (ARGs) and AMGs to determine whether *BL.*-derived proviral elements also employ synergistic evolutionary strategies. To start, we inspected prophage genomes for the presence of coding sequences (CDS) belonging to the Mo-AMG-HT PHROG category^73^. Of the 286 predicted prophages, 155 (∼54%) encoded 210 proteins assigned to this category. Notably, 93 of these proteins (∼44%) were functionally predicted to be associated with toxin-antitoxin systems. The presence of such proteins, which included Doc-like toxins, may enhance the vertical stability of prophages by functioning as addiction modules within their host^74,75,76^. Two widespread prophages also encoded the ‘ribonuclease toxin of AT system’ and were not targeted by any CRISPR spacers, highlighting that this toxin-antitoxin gene contributes to tolerance and maintenance within the host. Previous studies have also demonstrated that toxin-antitoxin systems can function to exclude competing MGEs, rather than acting strictly for stability^77,78,79,80^. We also detected 17 glutamine amidotransferase proteins, which are predicted to block superinfection of competing phages via cell wall‒ modification^81,82^. We further identified four specific superinfection exclusion proteins that function either to prevent receptor-binding^83,84^ or by blocking injection of viral DNA across the host cell membrane^85^. Overall, we find that *BL.*-derived prophages utilise multiple antagonistic strategies to enhance their persistence and maintenance within hosts, including metabolic modulation and “selfish” mechanisms that reduce competition from related phages.

We also screened prophages for ARGs and AMGs using a more targeted detection workflow to assess mutualistic bacteria‒phage interactions. Only one prophage identified in *BL. nexti* was predicted to encode ARGs. These ARGs included a major facilitator superfamily transporter of the DHA1 family associated with tetracycline resistance, a TetR/AcrR family transcriptional regulator (tetracycline repressor), and a VanZ-like family protein. The same prophage was also predicted to encode two potential AMGs: 3-oxoadipate enol-lactonase and S-adenosylmethionine (SAM) synthetase. 3-oxoadipate enol-lactonase is an enzyme critical for phthalate degradation that has been described in soil phages^86^. SAM synthetase has not been reported in viruses to date, but it is known to be involved in catalysing the formation of SAM, a methyl donor crucial for bacterial metabolism, and to possesses roles in bacterial immunity^87,88^.

Further analysis of potential AMG content revealed that the most prevalent protein was phosphoadenosine phosphosulphate (PAPS) reductase, which was identified in 37 prophages and located between viral genes on both sides in 21 of these (∼57%) sequences (viral-context inspection). PAPS reductase is involved in sulphur metabolism, catalysing a key step in the assimilation of sulphate into cysteine^89,90^. It converts 3ʹ-phosphoadenosine-5ʹ-phosphosulphate into sulphite, producing 3ʹ-phosphoadenosine-5ʹ-phosphate as a byproduct. Given that bifidobacteria generally lack genes encoding sulphur metabolism^90^, they cannot produce cysteine intracellularly and predominantly rely on acquiring it from the external environment^91^. It is therefore plausible that host strains utilise integrated proviruses with PAPS reductase to support sulphur assimilation. Indeed, PAPS reductase was present in a widespread *BL. infantis* prophage targeted by just three CRISPR spacers, indicating a high level of tolerance among host strains. However, no downstream genes related to the sulphur assimilation pathway were found in any of the strains with PAPS reductase that passed the viral-context inspection. Given that the product of PAPS reductase—sulphite—is reportedly toxic^92,93^, its presence is puzzling. However, we found sulphite exporters of the TauE/SafE/YfcA family in 19/21 (∼90%) of these prophage-harbouring strains, suggesting they can eflux toxic sulphite. The presence of PAPS reductase could therefore signify a cross-feeding function for other microbiota members by increasing the availability of a reduced sulphur source in the gut environment.

Given the uncertainty surrounding the presence and role of phage-encoded PAPS reductase in *BL.* strains, we clustered the corresponding protein sequences with homologues from UniProt (see Methods) to validate the PAPS reductase annotation (Figure S6). The PAPS reductase proteins we identified formed a distinct cluster with just four existing UniProt entries. All four are *Bifidobacterium*-derived PAPS reductase from alternative whole-genome sequencing datasets (UniProt accessions: A0A6A2S8H1, A0A6I2T478, A0A8U0LKI8, A0A9E7YXD8). For further validation of our PAPS reductase annotations, we predicted the protein structure of the cluster representative from our dataset using AlphaFold3^94^ and used Foldseek^95^ to compare it to the only available experimentally solved PAPS reductase crystal structure, which was from *Escherichia coli* (UniProt ID: P17854; PDB ID: 1SUR). The resulting model showed high global confidence (pTM = 0.92), demonstrating that the two proteins were structurally similar and likely share a similar overall function, supporting the prediction that *BL.* prophage-encoded PAPS reductase is involved in sulphur metabolism (Figure S7). In summary, the presence of potential AMGs, such as PAPS reductase, and ARGs suggests that certain *BL.* prophages do employ synergistic strategies that could enhance their persistence by improving host fitness.

### Prophages and CRISPR spacer arrays in *BL.* strains are highly stable in the first year of life

To evaluate how prophages and CRISPR spacer arrays change over time in the first year of life, we selected infant-derived *BL.* strains based on their presence at three or more timepoints (W2, M1, M2, M3, M6, M9 and M12) in samples taken from the same infants. Twenty-one strains from 21 infants met this criteria, comprising *BL. infantis* (n = 17), *BL. nexti* (n = 3), and *BL. suis* (n = 1). Among these ‘persistent’ bacterial strains, 17 harboured at least one prophage and six encoded CRISPR-Cas systems, with one genome containing two systems (*BL. nexti*, types I-E and II-C). The CRISPR-Cas systems from longitudinal strains included CRISPR-Cas type I-C (n = 3), type I-E (n = 3), and type II-C (n = 1). Two strains harboured neither a prophage or Cas system.

Investigations of the stability of longitudinal CRISPR spacer arrays revealed very minimal changes in array lengths across multiple timepoints. Only one strain showed an increase in spacer count—just a single spacer acquisition (*BL. infantis*, type I-C). We also observed a reduction in spacers in just a single strain (*BL. suis*, type I-E), with the number of spacers dropping from 52 spacers at M1 to 25 at M3. In the remaining four longitudinal strains with CRISPR-Cas systems, the spacer counts remained unchanged across all timepoints. To determine whether this stability also reflected sequence conservation, we compared spacer sequences and their order within individual strains across multiple timepoints. Nearly all strains displayed identical spacer content and arrangement (Figure 5a), with just a small subset of arrays showing limited spacer acquisition and rearrangement (Figure 5b). However, there was a considerable reduction in the size of the CRISPR array associated with a type I-E system present in a *BL. suis* strain, where there were no shared spacers between M1 and M2‒M3, as well as an overall reduction of 27 spacers (Figure 5c). As the arrays from M2 and M3 remained highly similar, this likely represents a strain-replacement event before M2, rather than a strong selective pressure that caused complete replacement and reformation of the CRISPR array.

**Figure 5.**
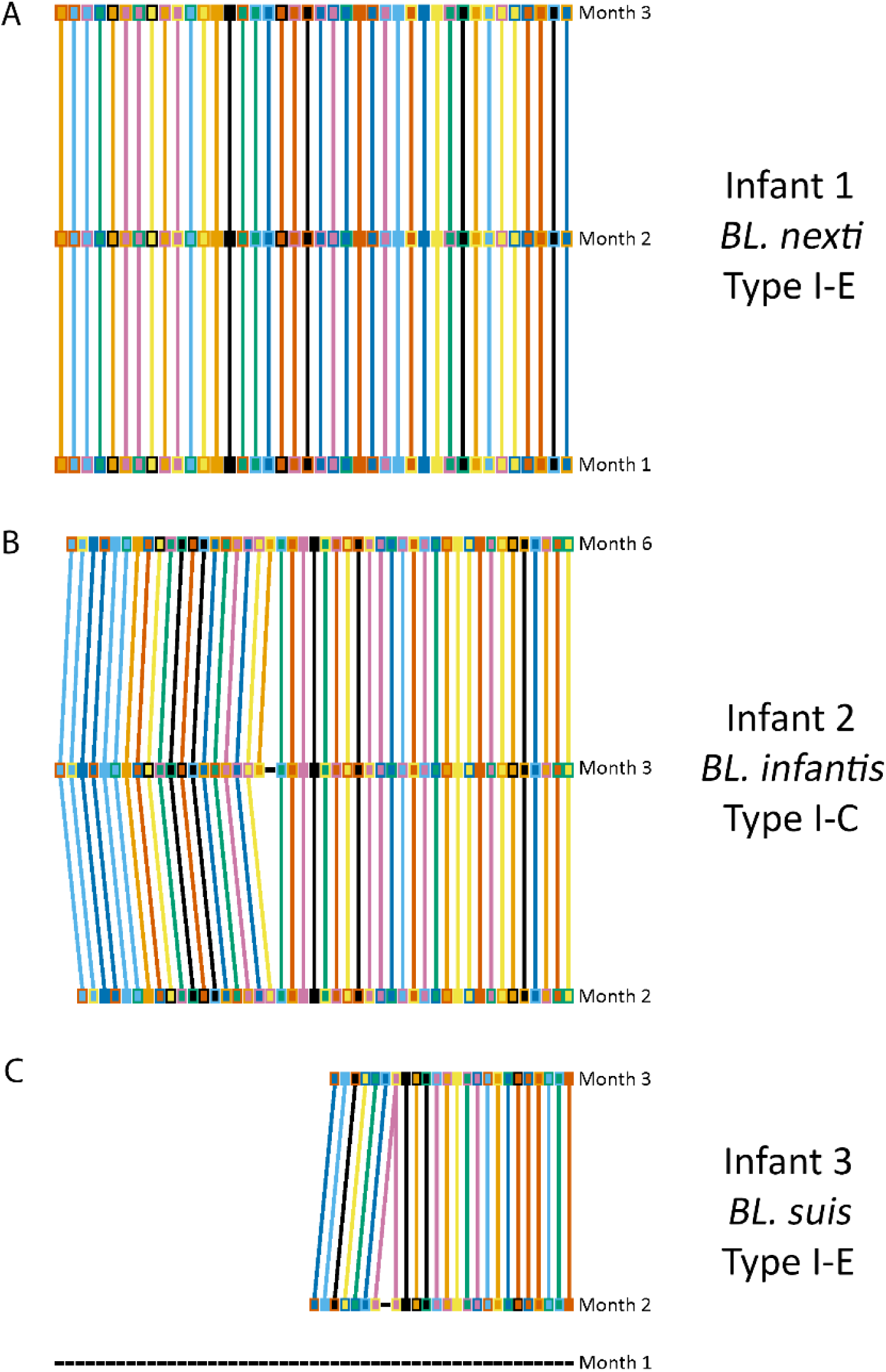
Longitudinal comparison of representative CRISPR spacer arrays within individual *BL.* strains. Unique spacers are represented by distinct combinations of fill and outline colours within panels. Spacers shared between timepoints are connected by lines. Dashed lines represent spacers with no shared sequences between timepoints **a**, Highly stable CRISPR spacer array from a Type I-E system (*BL. nexti*), with no changes observed across three months. **b**, CRISPR array from a Type I-C system (*BL. infantis*) showing the acquisition of two spacers at Month 3, whereby a single spacer was retained at Month 6. **c**, Example of a likely strain replacement event (*BL. suis*), where extensive spacer loss occurred between Months 1 and 2, followed by a stable but distinct array at Month 3.

Prophages also demonstrated remarkable stability across timepoints, showing high genomic conservation and very few signs of degradation. In three representative longitudinal *BL.* strains harbouring just a single prophage, including *BL. nexti* (Figure 6a) and *BL. infantis* (Figures 6b,c), viral sequences shared almost identical nucleotide identity at each time point, and their genome sizes remained unchanged. Across the entire sampling period, there were also no signs of gene loss/gain events or of recombination of viral sequences. This high level of conservation is indicative for prophages being stably maintained, with only limited prophage induction events taking place in this cohort of healthy infants.

**Figure 6.**
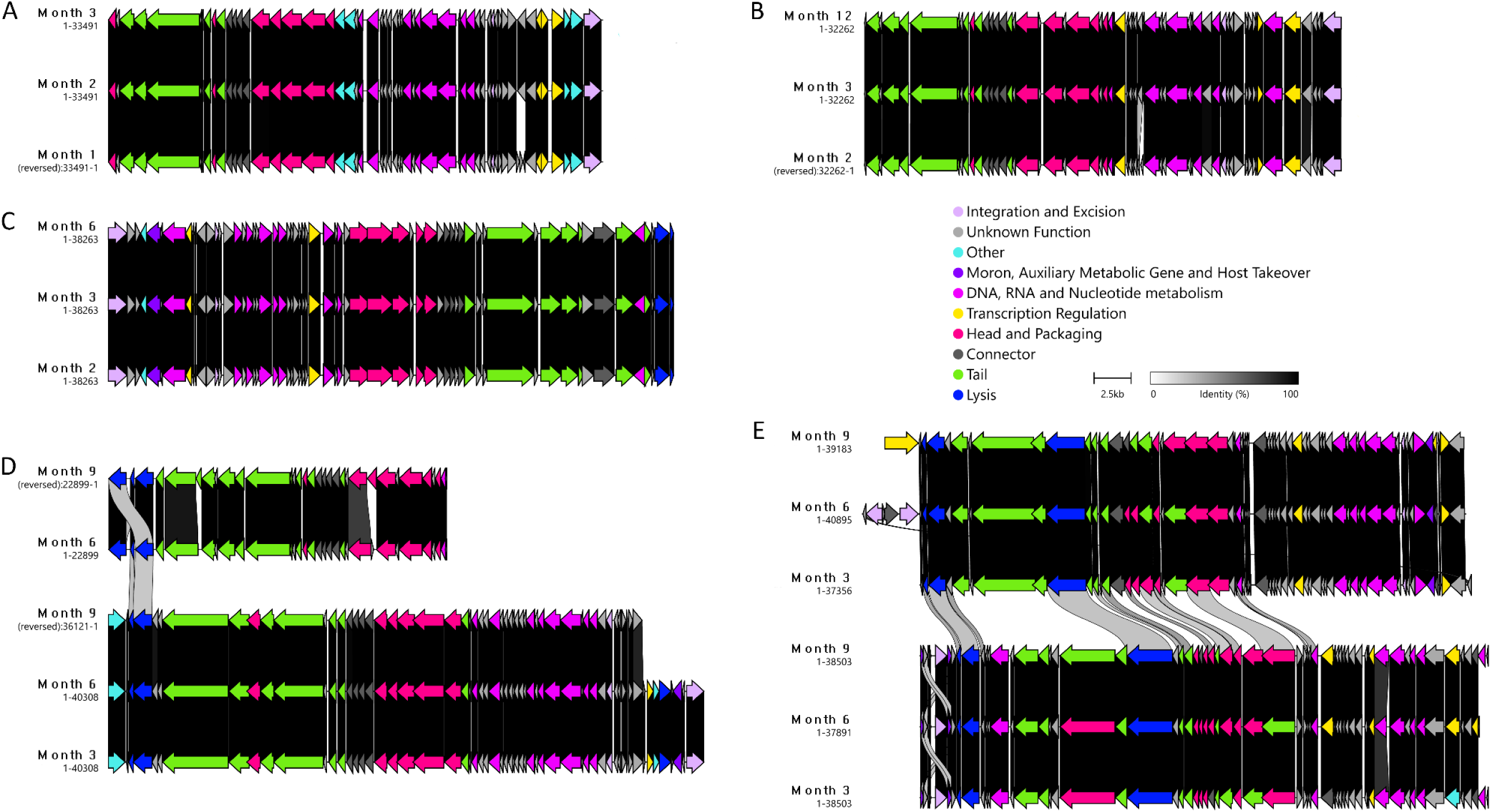
Longitudinal comparison of prophage sequences identified in *BL.* strains. Proviral integrity was largely maintained across three longitudinal time points in **a,** *BL. nexti* and **b,c,** *BL. infantis* strains, each harbouring just a single prophage. Prophages were also highly stable over time in *BL. infantis* strains containing two prophages (**d,e**), showing minimal changes to core viral genes. In panel **d,** there is a prophage integration event at Month 6, accompanied by reduced sequence identity of the major capsid protein at Month 9.

In strains harbouring multiple prophages, the prophages were also highly stable, showing very few changes in the core viral genome (Figures 6d,e). We also observed intermediate homology between co-resident prophages in both instances, with sequence identities of 30‒ 50% observed between several structural proteins, as well as amongst genes involved in nucleotide metabolism and lysis. Interestingly, we observed a potential new prophage infection event within the *BL. infantis* strain between M3 and M6 that was stably maintained through M9 (Figure 6d). However, there was a reduction in sequence identity observed between the major capsid proteins from M6 to M9.

Collectively, these results highlight the stability and tolerance of *BL.*-derived prophages, whereby virus‒host relationships are stably maintained in the infant gut and active prophage turnover is uncommon.

## Discussion

In this study we investigated if the prophages of infant-derived *Bifidobacterium longum* subspecies in a healthy Dutch cohort from Lifelines NEXT^36^ harness antagonistic and mutualistic evolutionary mechanisms to maintain their infectivity and persistence. Our data showed the presence of genomically diverse prophages in *BL.* strains, which were targeted by phage-specific defence mechanisms in the host. In response, these prophages have acquired specific anti-defence mechanisms to circumvent host defences. Despite these findings, prophages and CRISPR spacer arrays were highly stable throughout the first year of life. Prophages were also found to encode potential AMGs and other metabolism genes with the capacity to modulate host metabolic capacity.

This study highlights the genomic diversity, unique evolutionary trajectories, and several strategies employed by prophages of *BL.* strains that support their long-term maintenance and infectivity. Evidence of historical genomic recombination, horizontal gene acquisition, and accumulation of multiple anti-defence mechanisms demonstrate that these prophages utilise a range of antagonistic strategies to circumvent host immunity and potentially expand their host range. The presence of PAPS reductase and other metabolic genes in multiple prophages suggests that they may also engage in synergistic interactions with their host by enhancing host metabolic capacity and fitness, which promotes phage persistence by default. Contrary to recent reports of high dynamism within the infant gut virome^96,97^ and high turnover of CRISPR sequences in early life^98,99^, the integrated proviral sequences and CRISPR arrays of *BL.* strains were remarkably stable in this cohort of healthy infants. This stability points to a well-tolerated and evolutionarily stable relationship between *BL.*-derived prophages and their hosts during early life.

As human-associated *Bifidobacterium* prophages are yet to be isolated in the lab by means of viral plaque-forming units^45,46^, there are still large gaps in our knowledge of their biology and potential influence on human and infant health. To date, virulent bifidophages have been isolated only once for several phages infecting *Bifidobacterium asteroides* from the honeybee gut^100^. While recent works have highlighted the potential of bifidoprophages to drive bacterial shifts in the infant gut^44^, and with the genomic diversity and anti-defence system repertoire of these elements recently explored^41,42^, longitudinal analysis of prophage evolution specifically in *BL.* strains remains scarce. Here, we started filling this knowledge gap and unravelling these complex interactions. Future work could focus on real-time experiments in infant gut model systems seeded with natural infant microbial communities, which include *BL.* strains, to track prophage stability, CRISPR array turnover, and phage host-switching or successional phage dynamics. Additional research could also aim to experimentally validate anti-CRISPR activity and elucidate the functional impact of phage-encoded PAPS reductase to determine the ecological and evolutionary impact of these elements.

## Methods

### Sample collection, metagenomic sequencing, MAG assembly, and taxonomic classification

For this study, a total of 322 MAGs were selected for analysis, following the application of specific filtration criteria (as described below). Following dereplication at strain level, the final database of *BL.* strains included 213 MAGs from 139 infants from 137 families, comprising *BL. infantis* (n = 123), *BL. longum* (n = 53), *BL. nexti* (n = 31) and *BL. suis* (n = 6). All phenotypic and associated metadata for bacterial MAGs and prophages are provided in Supplementary Tables 1 and 2, respectively. Abbreviations used within these tables are defined in the accompanying “Abbreviations” sheet in each file.

For detailed information regarding sample collection and processing of faecal samples, as well as DNA extractions, sequencing, binning and MAG generation of infant samples from the Lifelines NEXT cohort study^36^, please see Sinha *et al*. (2024)^5^. Details regarding subspecies-level clustering of *BL. infantis*, *BL. longum*, *BL. nexti*, and *BL. suis*, are provided in Ennis *et al*. (2025)^35^.

### Filtering and dereplication of *B. longum* subspecies MAGs

Only high-quality MAG assemblies with completeness ≥90%, contamination ≤5%, and an N50 score ≥50,000, as determined by CheckM (v1.0.12; default within metaWRAP v1.3.2)^101,102^, were included in this study. Bacterial assemblies were dereplicated at strain level (99.9% average nucleotide identity (ANI)) using dRep (v3.4.2)^103^.

### Identification of CRISPR-Cas systems and spacer array analysis

CRISPRCasTyper (v1.8.0)^104^ was used to identify CRISPR-Cas systems and their associated spacer arrays in *BL.* genomes. Phage‒host relationships were predicted using the ‘*predictmatch*’ utility in SpacePHARER (v5.c2e680a)^105^. To detect unique spacer sequences, spacers were clustered using a sequence identity threshold of 100% with CD-HIT (v4.8.1)^106,107^.

### Detection of “high-quality” *BL.* prophages in dereplicated genomes

Putative proviral regions were detected in dereplicated bacterial MAGs using the ‘*end-to-end*’ command from geNomad (v1.11.0)^108^, under default parameters. All viral sequences predicted by geNomad were assessed for quality and completeness using CheckV (v1.0.1)^109^. Where applicable, CheckV trimmed regions of bacterial contamination flanking prophage ends. Viral sequences assigned as “Low-Quality” or “Not-Determined” and fragments below 5 kbp and above 100 kbp were removed from the database prior to further quality checks. Only prophage sequences assigned as “Medium-Quality” (≥50% completeness) or higher by CheckV underwent downstream viral quality control.

Next, we used the ‘*annotation*’ module from the PHORAGER pipeline (v0.2.0-beta) (https://github.com/aponsero/PHORAGER), which employs CheckV (v1.0.1)^109^, Pharokka (v1.7.5)^110^, and PHOLD v0.2.0^111^, to ensure that only prophages with ≥3 structural proteins (PHROG categories: ‘Head and Packaging’, ‘Tail’ and ‘Connector’^73^), comprising ≥20% of total CDS, were included in the final prophage database for downstream analysis.

### Characterisation of prophages and taxonomic classification

Taxonomic assignment of prophages at class level was performed using the ‘*annotate*’ module from geNomad^108^. Taxonomic classification at lower taxonomic levels (genus and species) was attempted using taxMyPhage (v0.3.4)^112^.

Viral genome clustering was performed using BLAST (v2.13.0)^113^, in conjunction with supporting code from CheckV (https://bitbucket.org/berkeleylab/checkv/src/master/)^109^, to determine genus- and species-level clustering of prophages based on pairwise ANI. Using parameters recommended by the Minimum Information about an Uncultivated Virus Genome (MIUViG)^114^, a ≥95% ANI cutoff and ≥85% target coverage was used to cluster viral species, with a ≥70% ANI cutoff used for genus clustering. Intergenomic similarity among all prophages was calculated based on pairwise ANI using CheckV supporting code.

Reciprocal intergenomic similarity was established by calculating the sum of the query and target sequence values, which were then halved. These values were then used to produce a two-way hierarchical clustering heatmap with Heatmap3 (v1.1.9)^115^ and ComplexHeatmap (v2.22.0)^116^ in R (v4.4.3)^117^ and RStudio (v2025.05.1+513)^118^. Whole proteomic trees were independently generated for all prophages and species representatives with ViPTreeGen (v1.1.3)^119^, and visualised in iTol (v5)^120^. Viral gene clusters were directly compared and visualised with Clinker (v0.0.31)^121^, whereby individual proteins were assigned specific colours based on PHROG functional categories^73^.

### Identification of anti-CRISPRs and alternative anti-defence systems

Acr proteins were predicted using Acafinder (version as of July 1, 2025; ‘*--Virus*’ option; guilt-by-association approach)^122^. DefenseFinder (v2.0.0; ‘*--antidefensefinder*’ module)^50,123^ and dbAPIS (a database of anti-prokaryotic immune system genes)^124^ were also used for the detection of alternative anti-defence systems in prophages.

### Determination of dN/dS ratios for anti-CRISPRs

Initially, Acr proteins with a minimum count of 20 were selected for selection pressure analysis. Here, dN/dS ratios were calculated using the Nei-Gojobori method (Fisher’s exact test)^68^, which estimates synonymous and non-synonymous nucleotide substitution rates. This statistical analysis was performed using MEGA12^125^.

First, nucleotide and amino acid sequences for each individual Acr type were aligned with MAFFT (v7.525)^126^. Using SeqKit (v2.10.0)^127^, the minimum and maximum lengths for each sequence was determined. These were then padded to ensure they were of equal size (see supporting code). Protein alignments were mapped to nucleotide sequences using PAL2NAL perl supporting code (https://www.bork.embl.de/pal2nal/) (v14)^128^ All identical sequences were deduplicated prior to dN/dS ratio calculations.

### Detection of antimicrobial resistance and AMGs

All prophages were annotated using Pharokka (v1.7.5)^110^ and PHOLD v0.2.0^111^, and protein sequences were subsequently used as input for the protein annotation tool MetaCerberus (v1.4.0)^129^, with the use of all available databases within the tool (--hmm ALL). Annotated proteins were first screened for viral hallmark genes, defined as those meeting at least one of the following criteria: (i) the presence of literal virus-related keywords (“portal”, “terminase”, “spike”, “capsid”, “sheath”, “tail”, “virion”, “holin”, “base plate”, “baseplate”, “lysozyme”, “head”, “structural”, “phage”, or “vir”) followed by manual verification^130^ and/or (ii) a VL-score > 4, following the approach described by Zhou et al (2024)^131^.

Genes not classified as viral hallmark genes during the initial screening were then manually inspected to identify potential ARGs and AMGs. To confirm the presence of both ARGs and AMGs within the prophage region, each candidate gene was evaluated for its genomic context^132,133^. Genes were retained as confirmed AMRs/AMGs only if they were flanked on both sides by previously defined viral hallmark genes.

To complement AMG identification and assess the presence of additional AMG pathway-associated genes within the bacterial genome, bacterial MAGs were also annotated using MetaCerberus (v1.4.0). Protein sequences were predicted using Prodigal (implemented within MetaCerberus)^134^, and annotation was performed against all available databases (-- prodigal, --hmm ALL). Resulting annotations were manually screened for the proteins of interest.

For clustering of protein sequences, all the annotated PAPS reductases available in UniProtKB as of 26 August 2025 that matched the query “phosphoadenosine phosphosulfate reductase” AND (taxonomy_id:2) were downloaded and merged with the predicted PAPS reductase sequences from this study. The combined dataset was clustered using CD-HIT (v4.8.1) with default parameters (90% sequence identity threshold for proteins)^106,107^. The representative sequence of the cluster containing the prophage-encoded PAPS reductases identified in this study was used for structure prediction with AlphaFold3 (default settings)^94^. The resulting model was compared with the experimentally determined crystal structure of the *Escherichia coli* PAPS reductase CysH (UniProt ID: P17854; PDB ID: 1SUR) using Foldseeek (v.10.941cd33)^95^.

### Identification of DGRs

To identify DGRs in prophages, viral sequences were screened using a multi-step pipeline adapted from Roux *et al*.^61^. First, gene prediction was performed with Prodigal-gv (v2.11.0)^134^, and reverse transcriptases (RTs) were detected through HMM-based annotation using hmmsearch (v3.4)^135^. Next, local sequence similarity searches were performed with BLASTn (v2.15.0) to identify repeat regions proximal to the RTs^113^, applying the parameters - word_size 8 -dust no -gapopen 6 -gapextend 2. Finally, putative DGRs were defined based on the presence of characteristic repeat structures and RT length criteria. Candidates with adenine mutation frequencies below 75% were excluded.

### Analysis of longitudinal prophage and CRISPR spacer data

Longitudinal faecal samples were collected from infants at set timepoints post-birth, including week 2 (W2) and between months 1‒12 (M1, M2, M3, M6, M9 and M12). Only bacterial strains present at three or more timepoints per individual were selected for downstream analysis of longitudinal samples. Bacterial genomes were once again processed with geNomad (v1.11.0)^108^ to obtain prophage coordinates. For closer examination and manual validation of each prophage identification step within longitudinal datasets, CheckV (v1.0.1)^109^, Pharokka (v1.7.5)^110^, and PHOLD v0.2.0^111^ were each run independently. Prophage sequences were then inspected for the presence of ≥3 structural proteins, comprising ≥20% of total CDS, as previously described when using the PHORAGER pipeline. To assess prophage stability, viral gene clusters of phages identified within individual strains at multiple timepoints were directly compared using Clinker (v0.0.31)^121^. For the identification of CRISPR spacer arrays in bacterial strains at each timepoint, CRISPRCasTyper (v1.8.0)^104^ was once again used. Spacer arrays from different timepoints were visualised and compared using CRISPR Comparison Toolkit (v1.0.3)^136^.

### Data analysis and visualisation

All statistical analyses were performed using base R statistical package (v4.4.3)^117^. Graphs were visualised using R package ggplot2 (v3.5.1)^137^ in RStudio (v2025.05.1+513)^118^ unless stated otherwise.

## Data availability

All MAGs analysed in this study are deposited in the European Nucleotide Archive (https://www.ebi.ac.uk/ena/browser/home) under the study accession number: PRJEB108306.

## Code Availability

All supporting code is publicly available on GitHub (https://github.com/JDocherty67/Bif_longum_prophages_LLNEXT_supporting_code).

## Acknowledgements

The data used in this manuscript are provided by LLNEXT. The authors are grateful for the participation of all the parents and infants in LLNEXT. We thank the LLNEXT team and maternity care providers for recruiting participants, collecting material during and after childbirth and building and maintaining the cohort. We thank Kate McInture for editing the text. We also thank the Genomics Coordination Center and the Center for Information Technology of the University of Groningen for their support and for providing access to the Gearshift, Peregrine, and Habrok high-performance computing clusters. The Lifelines NEXT cohort study received funds from the University Medical Center Groningen Hereditary Metabolic Diseases Fund, Health∼Holland (Top Sector Life Sciences and Health), the EU, the Northern Netherlands Alliance (SNN), the provinces of Friesland and Groningen, the municipality of Groningen, Philips, and the Société des Produits Nestlé. A.Z. is supported by the Nederlandse Organisatie voor Wetenschappelijk Onderzoek-VIDI (NWO-VIDI) grant 016.178.056, and NWO-VICI grant VI.C.232.074, EU Horizon Europe Program grant INITIALISE (101094099), NWO KIC grant KICH1.LWV04.21.01and NWO Gravitation grant Exposome-NL (024.004.017). M.Y. is the Rosalind, Paul and Robin Berlin Faculty Development Chair in Perinatal Research and is supported by the Azrieli Foundation grant for faculty fellows.

## Author Information

### Correspondence

Alexandra Zhernakova & James A. D. Docherty

### Contributions

J.A.D.D.: writing, methodology, investigation, data curation, visualisation, and formal analysis. A.Z.: supervision, funding, project administration, conceptualisation, methodology, resources, and validation. M.Y.: project administration, resources, conceptualisation, methodology, and validation. N.K., A.F-P.: formal analysis, writing, methodology, and conceptualisation. T.S., D.E., S.A-S., Sa.Ga.: resources, methodology, conceptualisation, data curation, and validation. C.M., Y.K.J., Se.Ge.: methodology and conceptualisation. M.F.B-G.: resources.

## Ethics declarations

### Research involving human participants

There were no human participants involved directly in this investigation; however, all *BL.* MAGs were derived from infant stool samples collected during the Lifelines NEXT cohort study^36^. The Lifelines NEXT study was approved by the ethics committee of the University Medical Center Groningen, document number METC UMCG METc2015/600.

