## Supplementary Information for "Prophages of infant-derived *Bifidobacterium longum* subspecies employ antagonistic and synergistic strategies to persist in their host"

J.A.D. Docherty *et al*.

##### Lifelines NEXT cohort study authors and affiliations

Milla F. Brandão-Gois[1], Marcel Bruinenberg[2], Siobhan Brushett[1,3], Jackie A.M. Dekens[1,4], Sanzhima Garmaeva[1], Sanne J. Gordijn[5], Soesma A. Jankipersadsing[1], Ank de Jonge[6,7], Gerard H. Koppelman[8,9], Marlou L.A. de Kroon[3], Folkert Kuipers[8,10], Alexander M. Kurilshikov[1], Anouk Marsman[2], Lilian L. Peters[6,7], Jelmer R. Prins[5], Sijmen A. Reijneveld[3], Sicco Scherjon[5], Jan Sikkema[4], Trishla Sinha[1], Johanne E. Spreckels[1], Aline B. Sprikkelman[8,9], Morris A. Swertz[1], Henkjan J. Verkade[8], Cisca Wijmenga[1], Alexandra Zhernakova[1]

[1] Department of Genetics, University of Groningen and University Medical Center Groningen, Groningen, the Netherlands

[2] Lifelines Cohort Study and Biobank, Groningen, the Netherlands

[3] Department of Health Sciences, University of Groningen and University Medical Center Groningen, Groningen, the Netherlands

[4] Innovation Center, University Medical Center Groningen, Groningen, the Netherlands

[5] Department of Obstetrics and Gynecology, University of Groningen and University Medical Center Groningen, Groningen, the Netherlands

[6] Department of Primary and Long-term Care, University of Groningen, University Medical Center Groningen, Groningen, the Netherlands

[7] Midwifery Science, Amsterdam University Medical Center, Vrije Universiteit Amsterdam, AVAG, Amsterdam Public Health, Amsterdam, the Netherlands

[8] Department of Pediatrics, University of Groningen and University Medical Center Groningen, Groningen, the Netherlands

[9] Groningen Research Institute for Asthma and COPD (GRIAC), University of Groningen and University Medical Center Groningen, Groningen, the Netherlands

[10] European Research Institute for the Biology of Ageing (ERIBA), University of Groningen and University Medical Center Groningen, Groningen, the Netherlands

### Supplementary Figures

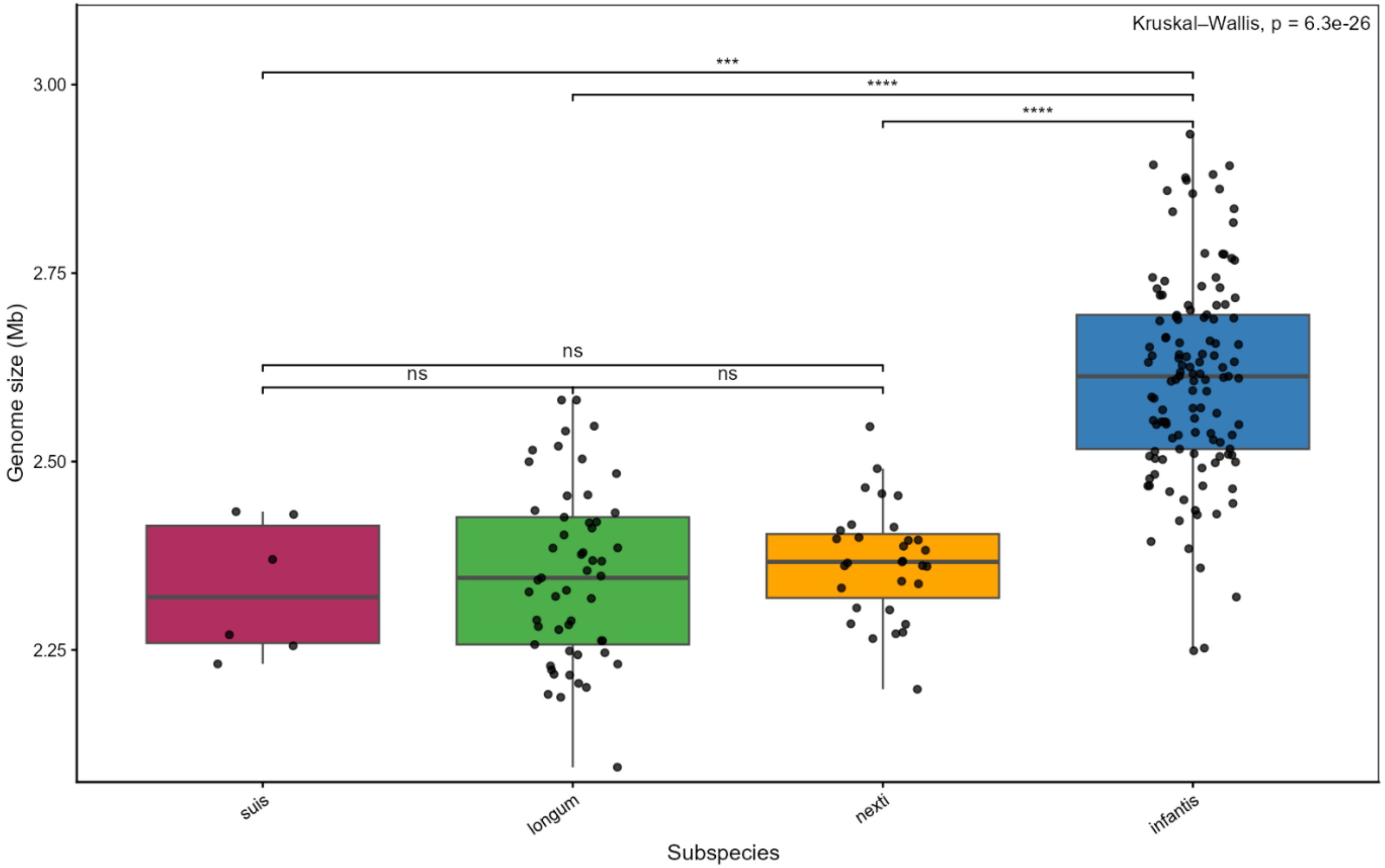

#### Figure S1. Genome size variation across *B. longum* subspecies.

Genome sizes (Mb) of bacterial MAGs from each *B. longum* subspecies (*BL.*), including *BL. suis*, *BL. longum*, *BL. nexti*, and *BL. infantis*. Individual genome sizes are depicted by dots. A Kruskal-Wallis test revealed significant differences in genome size among subspecies ($p = 6.3e^{-26}$). *BL. infantis* genomes are significantly larger than those of the other three subspecies (**p < 0.001, ***p < 0.0001, Dunn's post hoc pairwise comparisons). The genome sizes of *BL. suis*, *BL. Longum*, and *BL. nexti* did not differ significantly from one another (ns).

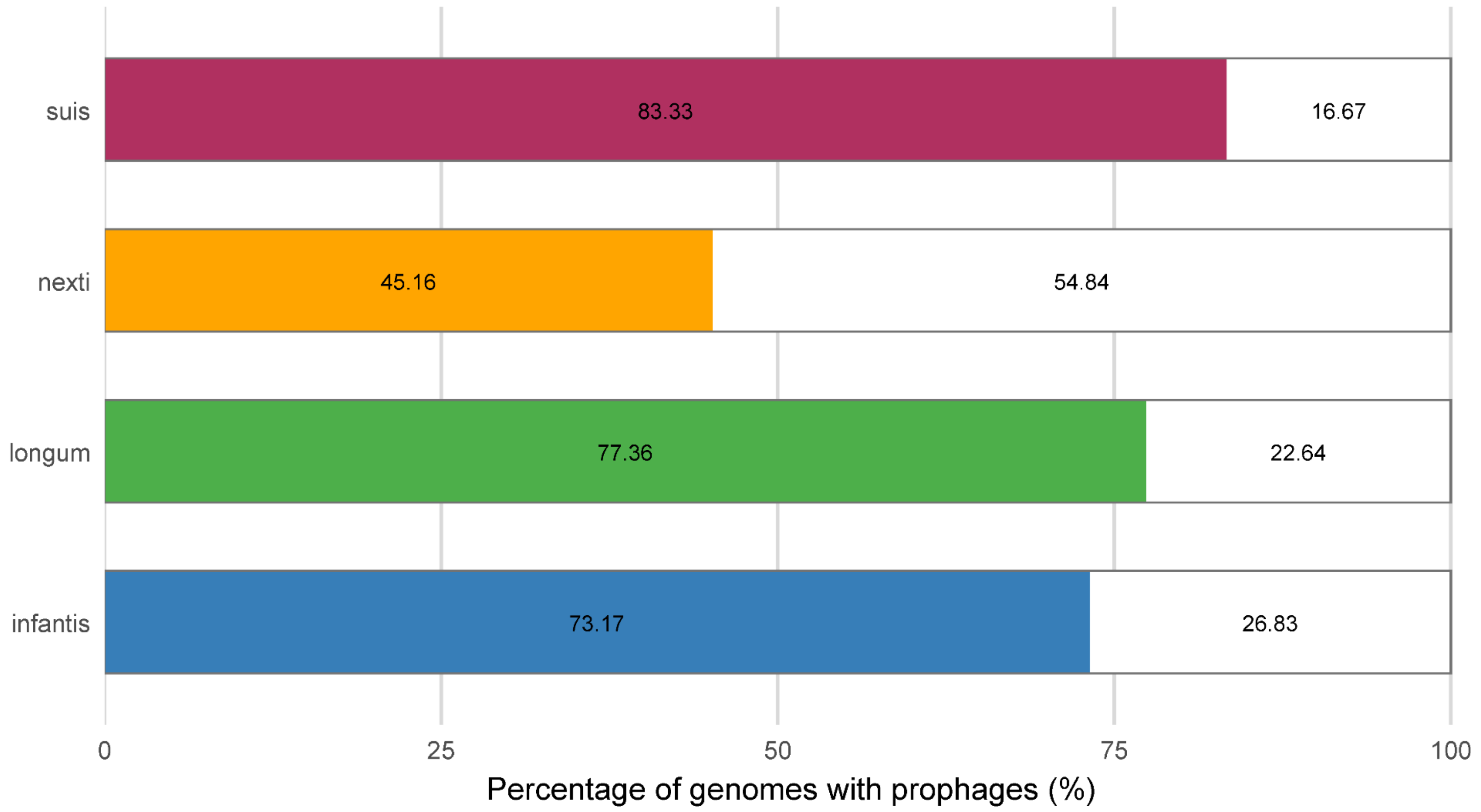

#### Figure S2. Prevalence of prophages in individual *BL.* subspecies.

Percentage of genomes within each *B. longum* subspecies that harbours at least one prophage (coloured portion) versus those lacking prophages (white portion). Values represent the proportion (%) of genomes harbouring prophages within each subspecies.

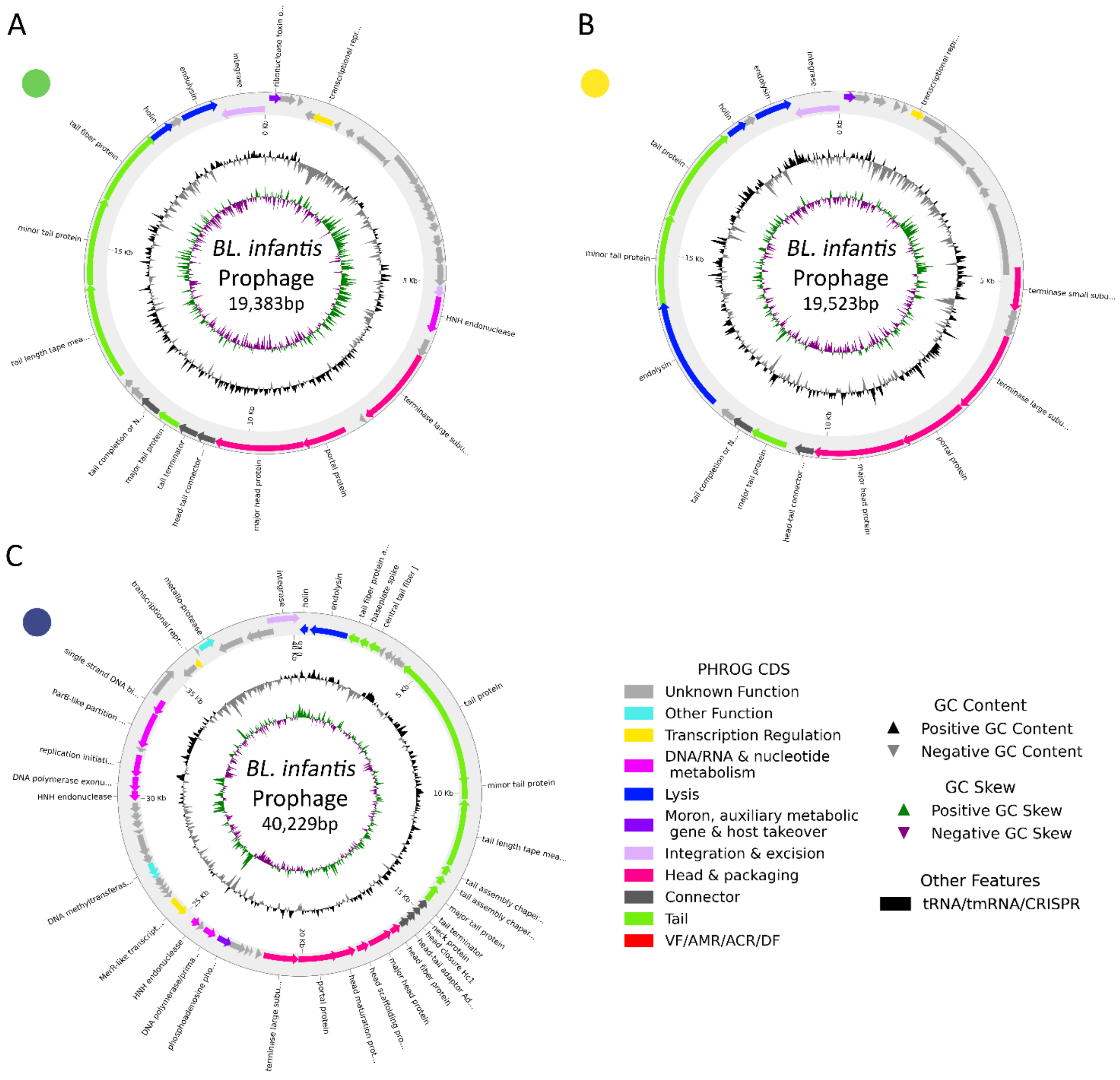

#### Figure S3. Circular genome maps of the three 'widespread' prophages that were each found to infect at least ten *BL. infantis* strains.

Circular representations of three *BL. infantis* prophage genomes, illustrating gene organisation, functional annotations, GC content, and GC skew. Coding sequences are coloured according to PHROG functional categories. The innermost rings display GC content (black) and GC skew (green/purple). tRNA/tmRNA/CRISPR features, when present, are indicated on the outer ring. **a-c**, Genome maps of three distinct prophages infecting *BL. infantis* with genome sizes of 19,383 bp (**a**), 19,523 bp (**b**), and 40,229 bp (**c**).

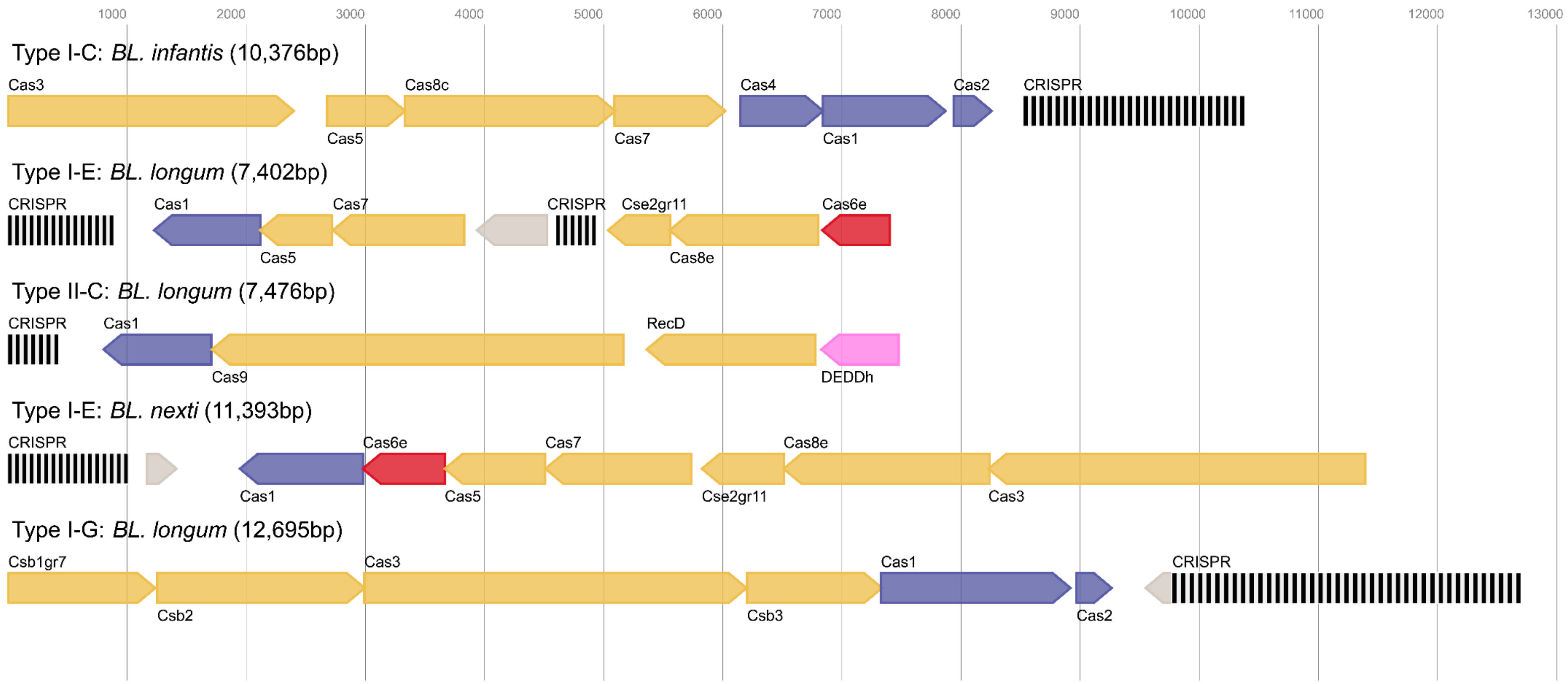

#### Figure S4. Complete gene maps of CRISPR-Cas systems detected in *BL.* strains.

Genomic organisation of representative CRISPR-Cas loci for the four subtypes identified in *BL.* strains, which included Types I-C, I-E, I-G, and II-C. The Type II-C system includes a degraded *cas*2 gene and encodes additional restorative proteins, including a DEDDh exonuclease and a RecD enzyme. A variant Type I-E locus was also identified, characterised by a truncated CRISPR array and substantial gene reorganisation.

CRISPR arrays are depicted by alternating black and white bars. Cas proteins involved in interference are shown in yellow, those involved in adaptation are shown in blue, and Cas6 ribonucleases are shown in red. Proteins of unknown function are marked in grey. The total locus size (bp) is indicated for each representative subtype.

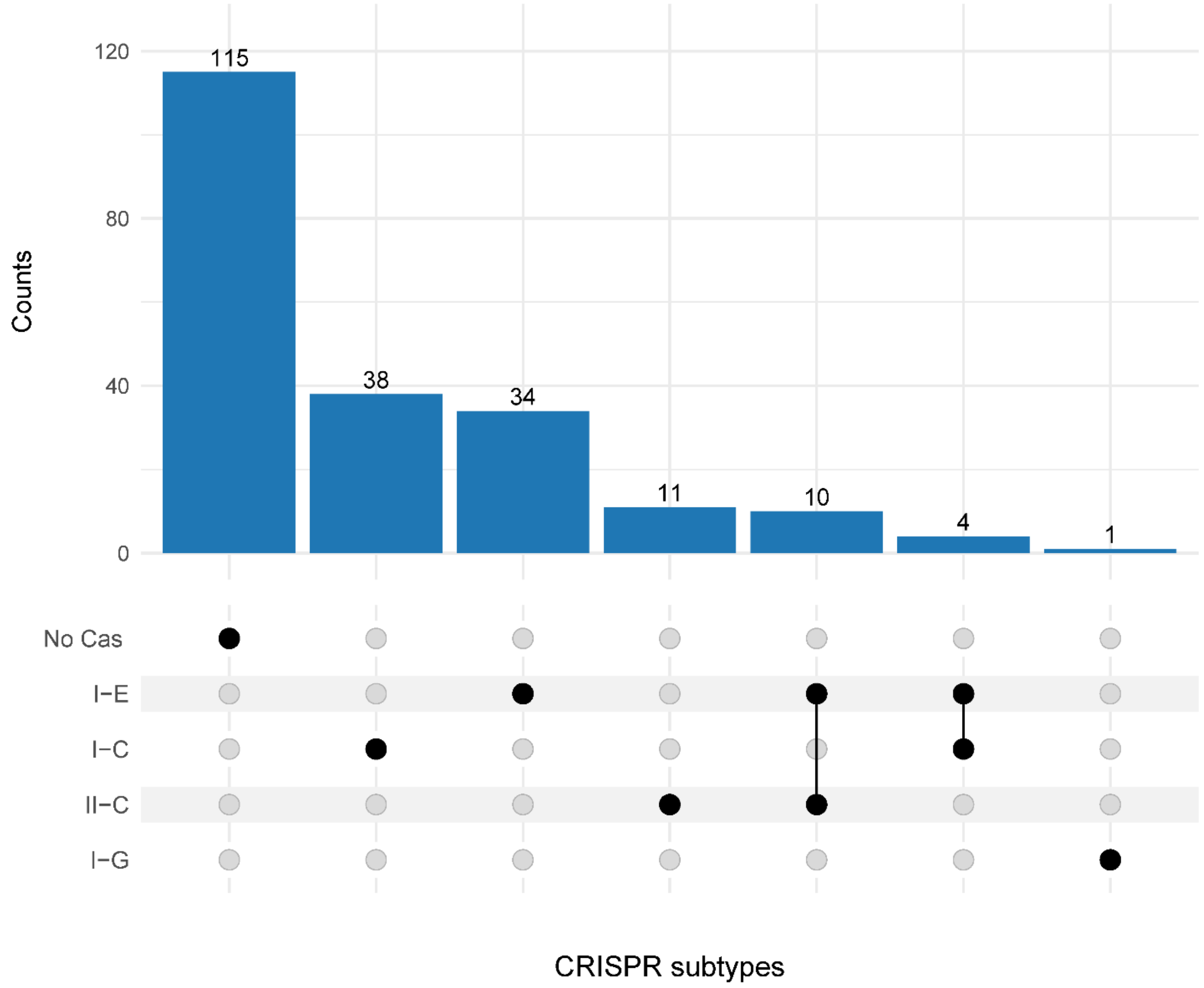

### Figure S5. Tallies of individual and co-occurring CRISPR-Cas systems in *BL.* strains.

Upset plot of the number of *BL.* strains with each distinct combination of CRISPR-Cas subtypes. In genomes with CRISPR-Cas systems, Types I-C and I-E dominated in the dataset. There were also 14 instances where there was a co-occurrence of multiple systems detected in individual genomes (I-E with II-C and I-C with I-E). Only one CRISPR-Cas Type I-G system was identified across all strains.

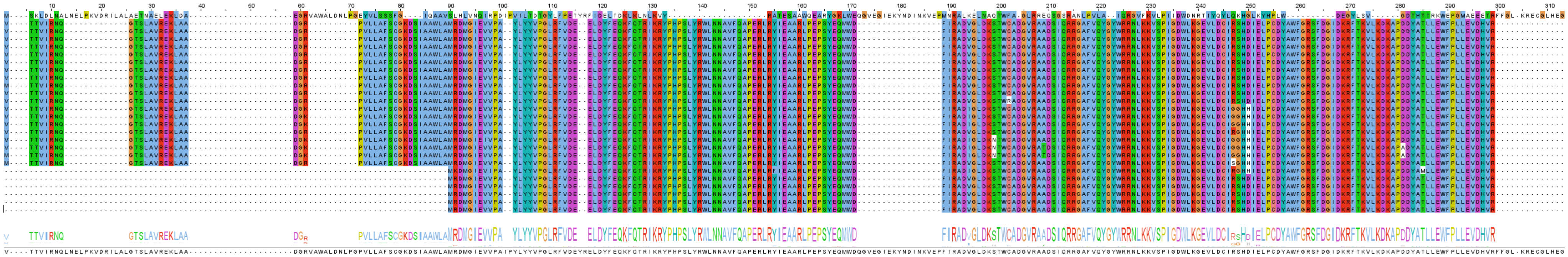

### Figure S6. Multiple sequence alignment of PAPS reductase protein sequences.

This alignment includes: (i) a reference PAPS reductase sequence from *Escherichia coli*, which served as the benchmark due to its experimentally confirmed enzymatic activity and solved crystal structure (line 1, ID: sp|P17854|CYSH_ECOLI) and (ii) sequences belonging to the PAPS reductase cluster with PAPS reductases predicted within *BL.* prophages in this study (lines 2–26). This set includes four PAPS reductase sequences from UniProt that were predicted in alternative *Bifidobacterium* species genomes from the other studies (UniProt IDs: ‘tr|A0A6A2S8H1|A0A6A2S8H1_BIFLN’, ‘tr|A0A6I2T478|A0A6I2T478_9BIFI’, ‘tr|A0A8U0LKI8|A0A8U0LKI8_BIFLI’, ‘tr|A0A9E7YXD8|A0A9E7YXD8_9BIFI’).

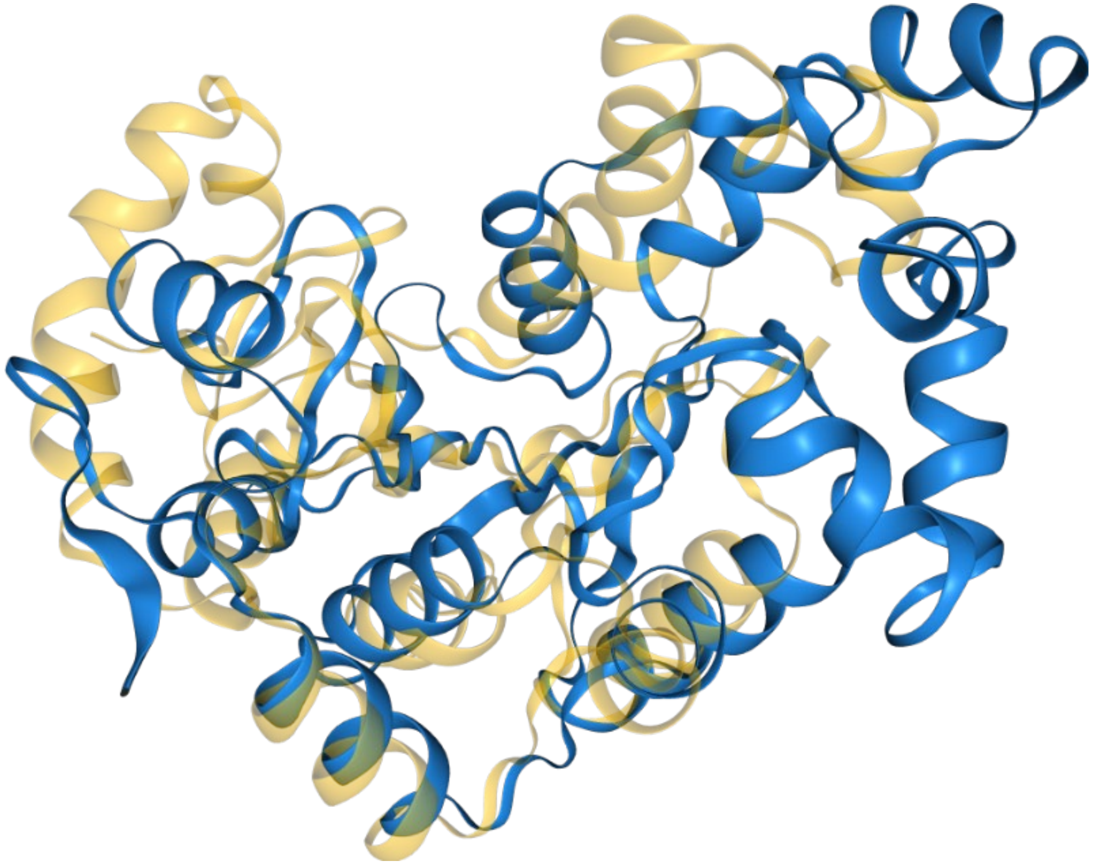

Figure S7. Superimposed ribbon diagrams of PAPS reductase from *BL.* prophage and *Escherichia coli*.

The blue ribbon represents the predicted AlphaFold model of the sequence representative belonging to the PAPS reductase cluster identified from *BL.* prophages in this study. The yellow ribbon represents the *E. coli* PAPS reductase (UniProt: P17854; PDB: 1SUR).
